# Psilocybin selectively rescues cognitive flexibility impairments caused by aberrant prefrontal error signaling

**DOI:** 10.64898/2026.07.05.736652

**Authors:** Christine Liu, Ellie M. Ho, Alexander S. Enos, Urte Beatrice Baublyte, Isabel M. Luna, Alexander R. Kosche, Vikaas S. Sohal

## Abstract

Psychedelic drugs show remarkable potential for treating psychiatric disorders, but the mechanisms underlying their therapeutic effects remain relatively unknown. Here, we demonstrate that psilocybin can powerfully ameliorate deficits in cognitive flexibility, but this effect depends on the specific circuit-level cause of those deficits. Using optogenetic models of cognitive inflexibility in mice, psilocybin rescued deficits caused by aberrant mesocortical signaling but failed to rescue deficits arising from disrupted interhemispheric gamma synchrony. Aberrant mesocortical signaling drove abnormally elevated activity in prefrontal cortex-mediodorsal thalamus (PFC-MD) projection neurons during post-error exploration, and psilocybin attenuated this pathological activity both acutely and 24 hours later. Patch-clamp electrophysiology revealed that psilocybin induces lasting plasticity in PFC-MD neurons, potentiating thalamic inputs while suppressing dopamine- and NMDA-receptor-dependent afterdepolarizations that could otherwise sustain aberrant post-error signaling. These findings reveal cellular and circuit mechanisms that could explain psilocybin’s therapeutic specificity and establish a precision medicine framework approach for psychedelic treatment.

## INTRODUCTION

Psychedelic drugs have shown immense potential for treating symptoms of psychiatric disorders that are characterized by maladaptive behaviors^1^, yet the associated mechanisms are unknown. A dominant hypothesis is that psychedelic drugs enhance cognitive and behavioral flexibility through facilitating neural plasticity^2^. Indeed, much evidence for structural plasticity^3,4^ and behavioral adaptation^5^ supports the notion that psychedelic drugs enable^6^ or restore^7^ enhanced learning. Yet, the effects of psychedelic drugs on cognitive flexibility have been variable^8^; some studies demonstrate positive outcomes^9,10^, no change^11^, or even worsened performance^12^. To maximize clinical impact, it is imperative to understand the sources of such variability, to clarify the suitability of psychedelic drugs for different psychiatric conditions.

Cognitive flexibility, the ability to adapt to changing environments, is impaired across depression, schizophrenia, addiction, Parkinson’s and other disorders^9,13,14^. Despite their distinct diagnoses and etiologies, people with these disorders often share difficulty in updating maladaptive thoughts and behaviors. Compared to healthy individuals, those with cognitive flexibility deficits struggle to adapt their behavior when the environment has changed, exhibiting poor performance in tasks such as the Wisconsin Card Sorting Task or Probabilistic Reversal Learning. Psychedelic drugs have been shown to improve cognitive flexibility in people with depression^10^ and Parkinson’s Disease^9^, but their effects on mood and cognition were not directly correlated. This suggests that deficits in cognitive flexibility, while present across many disorders, may be mediated by distinct neurobiological mechanisms from other symptoms and clinical features. In this context, animal models may be useful for identifying the specific mechanisms associated with deficits in cognitive flexibility and the therapeutic effects of psychedelics.

Our lab has characterized endogenous mechanisms for cognitive inflexibility in genetic mouse models^15^ and identified optogenetic manipulations that can induce impairments in otherwise healthy animals^16–19^. By leveraging this prior knowledge, we can test the effects of psilocybin on deficits caused by distinct circuit manipulations in the medial prefrontal cortex (mPFC) of mice. The mPFC is a critical node for cognitive flexibility, and thus manipulation of input and output projections can cause impairments. Our lab has previously shown that optogenetic stimulation of TH+ ventral tegmental area (VTA) inputs to the mPFC releases dopamine and glutamate and can impair cognitive flexibility in an odor-texture rule-shifting task^19^. Optogenetic inhibition of mPFC outputs to the mediodorsal thalamus (MD) impairs performance in a similar audiovisual rule-shifting task^18^. Impairing interhemispheric mPFC signaling also impairs cognitive flexibility, specifically when inhibiting callosal parvalbumin (cPV) projections during odor-texture rule shifts^17,20^. These circuit manipulations provide an opportunity to compare the therapeutic potential for psilocybin to rescue superficially similar deficits in cognitive flexibility that reflect distinct underlying circuit mechanisms.

## RESULTS

### Psilocybin alters strategy, but not performance, in a cognitive flexibility assay in otherwise-normal mice

As in prior studies^17,19–21^, we assessed cognitive flexibility using a digging-based, home-cage “rule-shifting” task adapted from attentional set-shifting assays. In this task (**Figure 1A**), mice learn an Initial Association (IA) based on a rule in one sensory dimension (i.e. texture) and must adapt when the rule is shifted to the other dimension (i.e. odor). To test the effects of psilocybin on cognitive flexibility, mice underwent one baseline day of testing before receiving an injection of either 2 mg/kg psilocybin or saline vehicle on Day 2. No significant differences in learning performance for the IA or Rule Shift (RS) were observed between saline and psilocybin (**Figure 1B**). A main effect of Day was observed for both drug conditions for IA performance, possibly due to increased familiarity with the task structure on Day 2. During the RS, mice encounter two different trial types: “non-conflict” trials, in which the correct choice is the same under both the IA and RS rules, and “conflict” trials, in which the IA and RS rules are in conflict (**Figure 1C**). Despite no overall difference in RS performance (**Figure 1B**), psilocybin-treated mice made significantly more non-conflict Errors on Day 2 (relative to the baseline day) compared to saline-treated controls (**Figure 1D**). Psilocybin did not significantly change the number of conflict errors made compared to saline controls (**Figure 1E**). How could psilocybin increase the number of non-conflict errors without worsening the overall performance of mice (i.e., the number of trials needed to reach the learning criterion)? To understand this, we computed how often mice exhibited ‘Win-Stay’ or ‘Lose-Shift’ behavior on non-conflict or conflict trials (based on the prior outcome from the same trial type) (**Figure 1F-N**). Consistent with the increase in non-conflict Errors, psilocybin significantly reduced non-conflict Win-Stay transitions (**Figure 1G**) meaning that mice were less likely to repeat correct choices on successive non-conflict trials. Psilocybin-treated mice also exhibited an increase in non-conflict Lose-Shift transitions (**Figure 1J**), revealing that they were also more likely to change to a correct choice after making an incorrect non-conflict choice. Together, these results show that in previously-normal mice, psilocybin promotes flexibility in decision-making during non-conflict trials. Psilocybin’s effects on learning strategy are unlikely to be caused by generic impairments in memory or learning, as mice in both drug conditions made similar total numbers of errors and proportionally more conflict errors, reflecting retention of the IA Rule (**Figure S1A-E**). The effects of psilocybin are preserved when examining each sex separately (**Figure S1F-P**), demonstrating the changes that acute psilocybin exerts on learning strategies as mice adapt to rule changes in the rule-shifting task affect males and females similarly.

**Figure 1.**
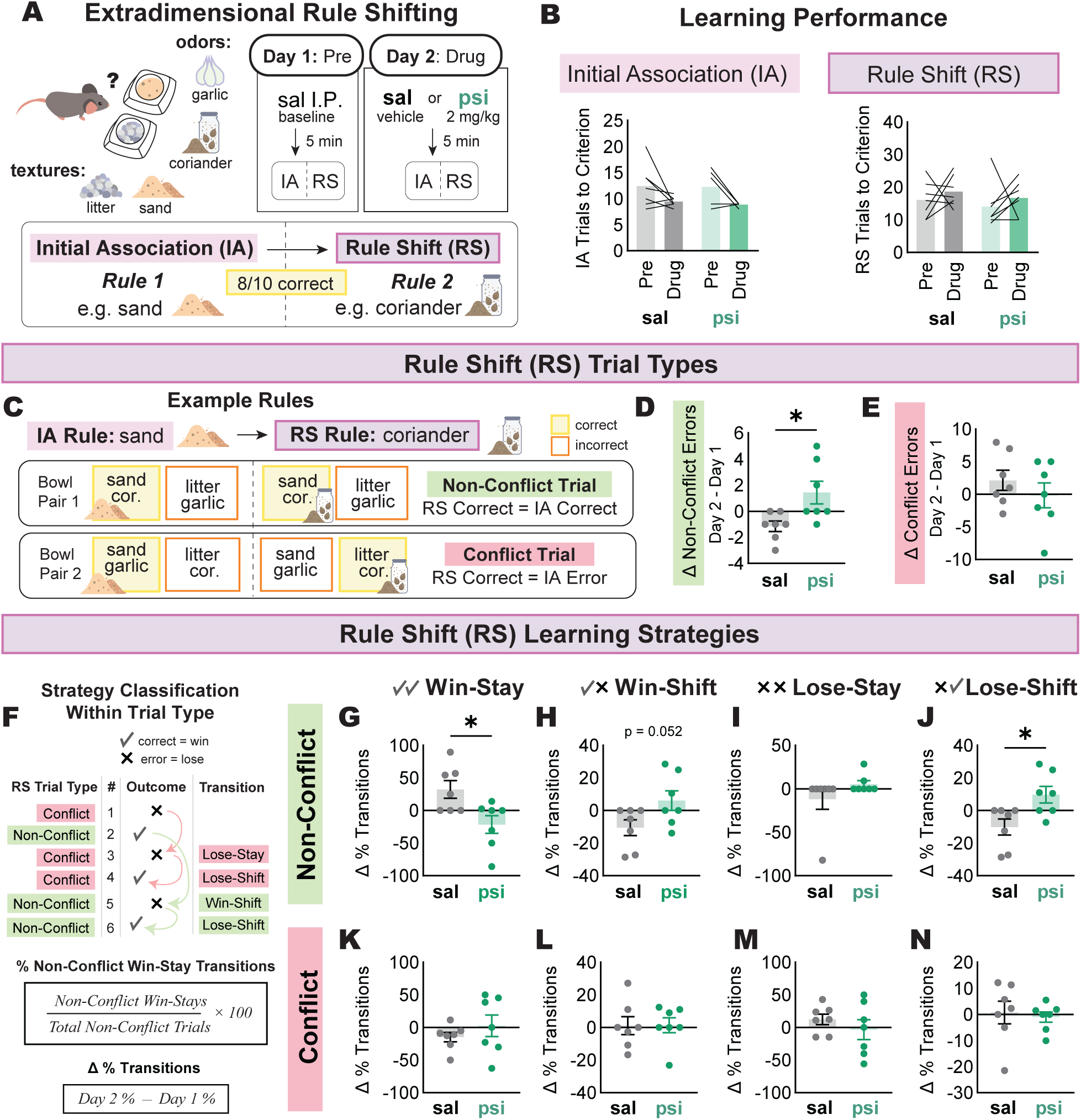
Psilocybin alters strategy, but not performance, in a cognitive flexibility assay in otherwise-normal mice. A. Schematic of home cage extradimensional rule shifting task and experimental timeline. B. Trials required to reach 8/10 criterion for the Initial Association (IA, left) and Rule Shift (RS, right) stages of the task. Bars represent means and lines represent individual animals. C. Schematic defining conflict and con-conflict Trials based on example IA and RS rules. D. Mice receiving acute psilocybin treatment on Day 2 (n = 7) exhibited more errors during non-conflict Trials (relative to the baseline day) compared to saline controls (n = 7). E. No difference in the number of conflict errors relative to baseline in animals receiving saline or psilocybin. F. Schematic for defining transitions within trial types and example equation for quantifying change in % transitions between days. G. Psilocybin treatment decreases % non-conflict Win-Stay transitions. H. Psilocybin trends toward an increase of non-conflict Win-Shift transitions. I. No difference was observed across days or drug treatment in non-conflict Lose-Stay transitions. J. Psilocybin increases % non-conflict Lose-Shift transitions compared to saline controls. K. No differences were observed for % conflict Win-Stay transitions based on drug treatment. L. Same as K, but for % conflict Win-Shift transitions. M. Same as K, but for % conflict Lose-Stay transitions N. Same as K, but for % conflict Lose-Shift transitions. *p < 0.05, by two-way repeated measures ANOVA (A; * main effect of day p = 0.01, B; NS) or unpaired two-tailed t-test (D-E, G-N). Bars represent means, error bars represent SEM, and dots represent individual animals (D, E, G-N). See also Figure S1.

### Psilocybin increases exploratory learning strategies and rescues cognitive flexibility impaired by stimulation of TH+ VTA → mPFC projections

Given the transformative potential for psilocybin to treat cognitive flexibility impairments in psychiatric disorders, we sought to evaluate the extent to which psilocybin could rescue deficits caused by aberrant mesocortical signaling. Prior work in the lab demonstrated that a single, unilateral burst of channelrhodopsin (ChR2) stimulation delivered to TH+ VTA → mPFC projections immediately following RS error choices was sufficient to impair performance in the rule-shifting task^19^. Indeed, we were able to replicate this effect when using either terminal stimulation in the mPFC (**Figure S2A-H**) or projection-specific cell body stimulation in the VTA (**Figure 2A, S2I**). To test whether psilocybin treatment could rescue this impairment in cognitive flexibility caused by aberrant mesocortical outcome signaling, animals underwent 3 days of rule-shifting (Day 1: no stimulation, Day 2+3: optogenetic stimulation) with counterbalanced drug treatment on Days 2 and 3. We compared the effects of optogenetic mesocortical stimulation, i.e., the change in performance compared to Day 1 (no stim), in mice that received psilocybin treatment acutely or 24 hours prior (**Figure 2B**). As expected, there were no differences observed in IA learning performance across Virus or Drug Order conditions and unilateral burst stimulation on RS errors impaired cognitive flexibility by increasing the number of trials to reach RS criterion (**Figure 2C-D**). Remarkably, acute psilocybin restored cognitive flexibility when administered prior to optogenetic stimulation, whether acutely or 24 hours prior (**Figure 2D**). Consistent with prior work^19^, TH+ VTA → mPFC burst stimulation causes an increase in both non-conflict and conflict Errors (**Figure 2E-F**). To our surprise, psilocybin treatment reduced errors across both trial types (**Figure 2E-F**). Examining learning strategies across trial types revealed changes caused by TH+ VTA → mPFC burst stimulation and the effects of psilocybin administered acutely or 24 hours prior (**Figure 2G-N**). Specifically, conflict Lose-Stay transitions are increased in ChR2 mice when receiving optogenetic stimulation but decreased by psilocybin (**Figure 2M**). There was a corresponding increase in conflict Lose-Shift transitions in the ChR2 mice which received psilocybin (**Figure 2N**). We also observed the expected effects of acute psilocybin in the eYFP mice that received saline first, namely a decrease in non-conflict Win-Stay transitions and increased non-conflict Lose-Shift transitions (**Figure 1G, 1J**). Of note, we did not observe these effects in the eYFP mice that received psilocybin first, suggesting that the effects of psilocybin in otherwise-normal mice are not always reliably detected, possibly due to the low number of trials (∼15) that unimpaired animals need to learn the RS (**Figure 1B**). In animals that are impaired by TH+ VTA → mPFC burst stimulation, we observe drastic changes in both RS performance and the underlying learning strategies. Together, these data demonstrate that psilocybin protects against impairments caused by TH+ VTA → mPFC burst stimulation, altering learning strategies and restoring performance to levels seen in eYFP controls in the absence of drug treatment.

**Figure 2.**
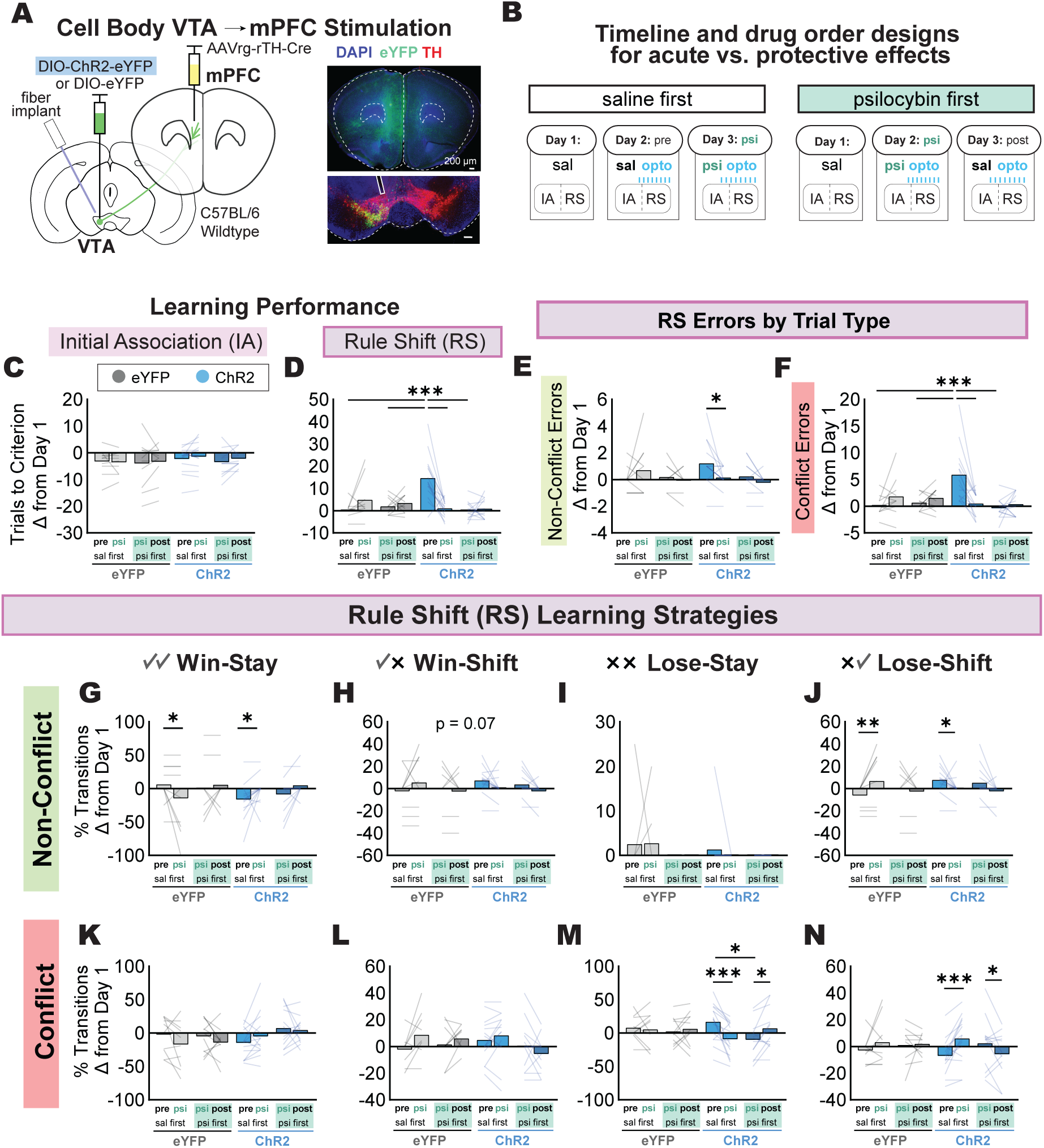
Psilocybin increases exploratory learning strategies and rescues cognitive flexibility impaired by stimulation of TH+ VTA → mPFC projections. A. Schematic for viral targeting of TH+ VTA neurons that project to mPFC with ChR2 (left). AAVrg-rTH-Cre was injected in the mPFC for retrograde Cre expression and AAV5-DIO-ChR2-eYFP or AAV5-DIO-eYFP control was injected in the ipsilateral VTA. An optical fiber implant was positioned above the VTA for cell body stimulation. Right: representative histological images for the mPFC (top) and VTA (bottom). Scale bar = 200 microns. B. Schematic for experimental timeline with 2 drug orders: “saline first” or “psilocybin first”. All mice underwent one baseline testing day (Day 1) with saline treatment and no optogenetic stimulation. On Days 2 and 3, all mice received a single unilateral burst of TH+ VTA → mPFC stimulation upon making an RS error. The drug groups differed based on whether they received saline first (Day 2: saline, Day 3: psilocybin) or psilocybin first (Day 2: psilocybin, Day 3: saline). C. No change in trials to reach criterion for the Initial Association (IA) relative to baseline were observed between groups based on Virus (eYFP or ChR2) or Drug Order (sal first or psi first). eYFP sal first n = 10, eYFP psi first n = 11, ChR2 sal first n = 15, ChR2 psi first n = 13. D. ChR2 mice that received saline first required significantly more trials to reach Rule Shift (RS) criterion on Day 2 compared to Day 3 and compared to all other groups on Day 2. E. Non-conflict errors were significantly increased in ChR2 mice that received saline first on Day 2 compared to when they received psilocybin on Day 3. F. ChR2 mice that received saline first made significantly more conflict errors on Day 2 compared to Day 3 and compared to all other groups on Day 2. G. eYFP mice that received saline first exhibited a decrease in % non-conflict Win-Stay transitions on Day 3 (psi) compared to Day 2 (sal). In contrast, ChR2 mice that received saline first had decreased % non-conflict Win-Stay transitions on Day 2 (sal) compared to Day 3 (psi). H. The change in % non-conflict Win-Shift transitions showed a trend-level interaction between Virus and Drug Order. I. No differences were observed for change in % Non-Conflict Lose-Stay transitions. J. eYFP mice that received saline first exhibited an increase in % non-conflict Lose-Shift transitions on Day 3 (psi) compared to Day 2 (sal). In contrast, ChR2 mice that received saline first had increased % non-conflict Lose-Shift transitions on Day 2 (sal) compared to Day 3 (psi). K. No differences were observed for the change in % conflict Win-Stay transitions. L. No differences were observed for the change in % conflict Win-Shift transitions. M. ChR2 mice that received saline first exhibited a significant increase in % conflict Lose-Stay transitions on Day 2 (sal) compared to Day 3 (psi). For ChR2 mice, there was also a significant increase in % conflict Lose-Stay transitions on Day 2 for saline first compared to psilocybin first. A significant difference was also observed between days in ChR2 mice that received psilocybin first, with decreased % conflict Lose-Stay transitions in acute psilocybin (Day 2) compared with 24 hours later (Day 3). N. Both ChR2 mice that received saline first and ChR2 mice that received psilocybin first exhibited significantly decreased % conflict Lose-Shift transitions on the days that they received saline compared to the days that they received acute psilocybin. *p < 0.05, ** p < 0.01, *** p < 0.001 by two-way repeated measures ANOVA (C-N) and Tukey’s post-hoc test (D, E, F, G, J, M, N). Bars represent means, error bars represent SEM, and dots represent individual animals. See also Figures S2 and S3.

It is worth noting that psilocybin failed to rescue superficially similar cognitive flexibility deficits when they were induced by a distinct neurobiological mechanism. We replicated prior work from the lab^17^ which optogenetically inhibited callosal parvalbumin projections (cPV), and found that this disrupts mPFC gamma synchrony and worsens RS performance, specifically increasing conflict errors (**Figure S3A-F**). Not only did acute psilocybin treatment fail to rescue RS performance, psilocybin increased non-conflict errors (**Figure S3G**). To examine underlying learning strategies, we quantified Win-Stay/Lose-Shift transitions (**Figure S3H-O**), but there was not a single clear factor that explained the optogenetically induced behavioral impairment in eNpHR mice. Deficits caused by cPV inhibition may be due to impaired attention or other cognitive domains that mPFC PV cells are known to mediate^16,22,23^. Acute psilocybin did appear to alter learning strategies by increasing non-conflict Win-Shift transitions in eNpHR mice (**Figure S3I**), explaining the increase in non-conflict errors (**Figure S3G**). There was also a main effect of Day observed for conflict Lose-Stay transitions (**Figure S3N**). When performing five consecutive days of rule-shifting, it is perhaps unsurprising that animals may alter their learning strategies over time. Related to this, we did not observe a change in learning strategies when unimpaired animals were treated with psilocybin on Day 4, and this may reflect the fact that the task was already quite familiar to animals by this point (in contrast to our prior experiments which gave psilocybin to animals that only performed 2-3 days of rule-shifting; **Figures 1 and 2**). Together, the effects of psilocybin in healthy and impaired animals suggest that psilocybin may alter learning strategies employed by animals in flexible decision-making. Whether psilocybin can reverse deficits in cognitive flexibility depends on the neurobiological mechanisms underlying that pathological state.

### Psilocybin attenuates abnormally elevated post-error activity in mPFC → MD neurons

Based on accumulating evidence that psilocybin alters learning strategies, we next explored the activity of prefrontal neurons that project to the mediodorsal (MD) thalamus, a population known to play key roles in cognitive flexibility and decision-making^18,21,24^. In addition to the viral injections and optical fiber implants required for cell body stimulation of TH+ VTA → mPFC projection neurons in wildtype mice, we injected AAVrg-FlpO in the MD thalamus and AAV-Coff/Fon-sRGECO in the mPFC bilaterally (**Figure 2A, S4A**). This strategy enabled recordings of calcium activity in mPFC → MD cells across 3 days of rule-shifting with counterbalanced drug order to test the acute and persistent effects of psilocybin on behavior and neural activity (**Figure 3B, S4B**). Psilocybin rescued cognitive flexibility both acutely and 24 hours after administration (**Figure 3C**), demonstrating a protective effect against impairments caused by TH+ VTA → mPFC stimulation. Given that impairments are caused by unilateral burst stimulations delivered upon RS errors, we sought to quantify ipsilateral neural activity both during the immediate response to errors and during post-error exploration (**Figure 3D**). On the baseline day, mPFC → MD activity is elevated during the immediate (0-5s) response to error outcomes, but not correct outcomes (**Figure S3E, S4C**). There was no difference across groups or days for this immediate response to errors (**Figure S4D**). However, when examining mPFC → MD activity during the 10-90s post-error period when animals engage in extended exploration of the remaining, incorrect bowl, we observed abnormally elevated activity in ChR2 mice that had not yet received psilocybin (**Figure 3F**). Consistent with the behavioral rescue we observed, both acute psilocybin and psilocybin treatment 24 hours prior attenuated this abnormally elevated mPFC → MD activity in ChR2 mice. Of note, persistent effects of psilocybin were also apparent when examining the immediate response (0-5s) after IA errors, with decreased activity in eYFP and ChR2 mice 24 hours after psilocybin administration (**Figure S4E**), despite no differences in the 10-90s post-error exploration period (**Figure S4F**). The effects of psilocybin on mPFC → MD activity were primarily observed after errors. No difference in activity was observed after IA correct outcomes and only a main effect of Group observed for activity after RS correct outcomes (**Figure S4G-J**). Together, these behavioral and neural data suggest that psilocybin has acute effects on mPFC → MD neuronal activity after RS errors and provides a protective effect even after the drug has washed out. Changes in neural activity that persist at least 24 hours after psilocybin treatment may reflect plasticity within this population itself, and/or of its inputs.

**Figure 3.**
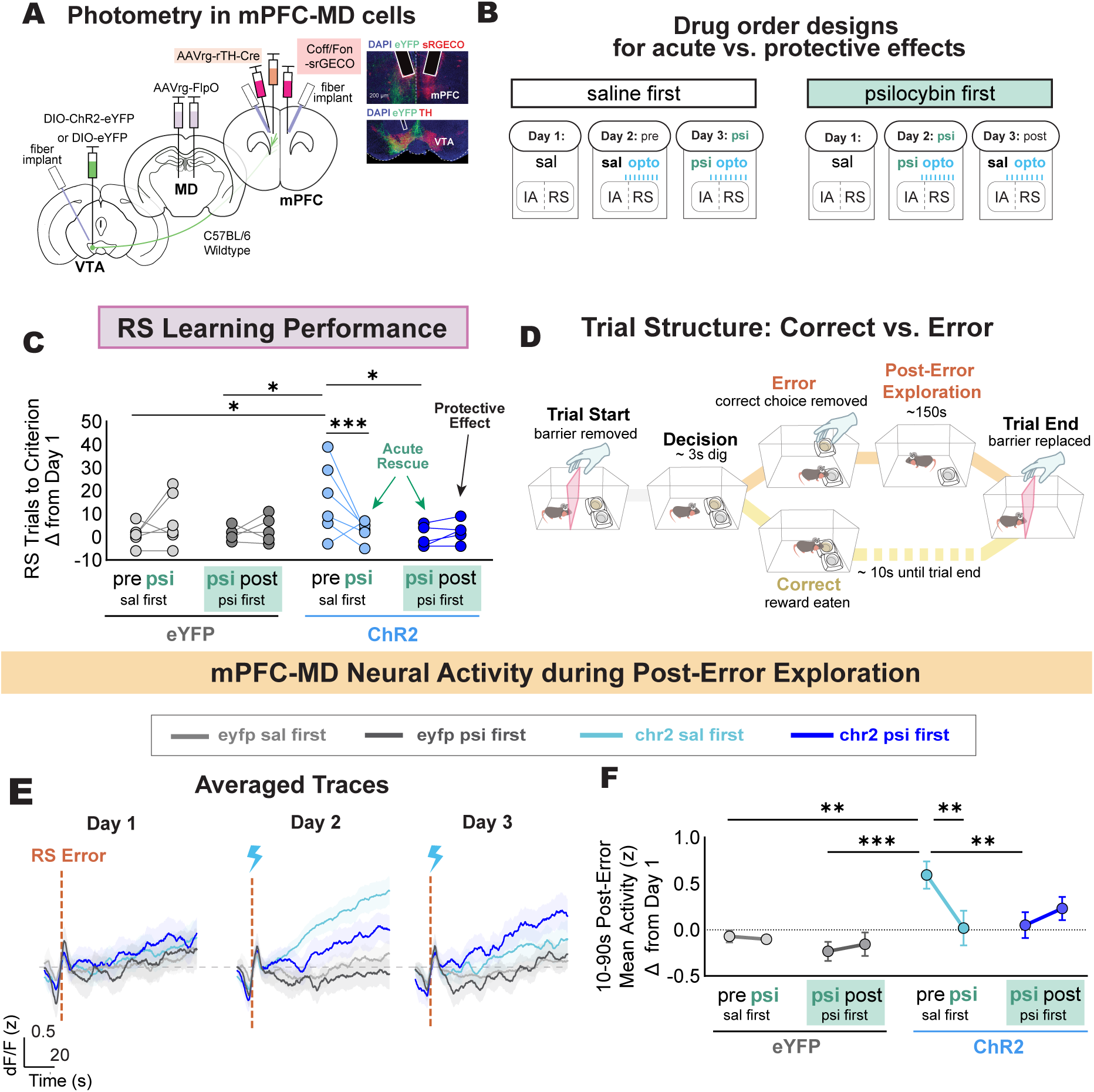
Psilocybin attenuates abnormally elevated post-error activity in mPFC → MD neurons. A. Schematic for cell body TH+ VTA → mPFC stimulation plus photometry to measure genetically-encoded calcium sensor signals from mPFC → MD cells (left). Wildtype mice are injected unilaterally with AAVrg-rTH-Cre in the mPFC for retrograde Cre expression in TH+ cells. AAV5-DIO-ChR2-eYFP (experimental group) or AAV5-DIO-eYFP (control) is injected ipsilaterally in the VTA. An optical fiber implant is positioned above the VTA for delivery of 473 nm laser stimulation. AAVrg-FlpO is injected bilaterally in the mediodorsal thalamus for retrograde Flp expression. AAV-CoFF/Fon-sRGECO is injected bilaterally in the mPFC and fiber implants are positioned above mPFC for photometry recordings. Right: representative histological images. B. Experimental timeline: Day 1 - all mice undergo a baseline day of testing with saline and no optogenetic stimulation. Mice in the “saline first” group receive saline and optogenetic stimulation on Day 2, then psilocybin and optogenetic stimulation on Day 3. Mice in the “psilocybin first” group receive psilocybin and optogenetic stimulation on Day 2, then are tested with saline and optogenetic stimulation on Day 3 (24 hours after psilocybin treatment). C. On Day 2, ChR2 saline first animals (n = 6) require significantly more trials to reach the RS learning criterion (relative to baseline) compared to both eYFP mice (sal first n = 6; psi first n = 5) and ChR2 mice that receive psilocybin first (n = 5). On Day 3, acute psilocybin treatment restores normal performance in ChR2 saline first animals. ChR2 animals that received psilocybin first on Day 2 continue to exhibit normal RS performance on Day 3 (24 hours post psilocybin treatment). D. Schematic of trial structure for incorrect (top) and correct (bottom) trials. When animals make an incorrect choice, the bowl containing the reward is removed by the experimenter but the animal can explore the cage and the remaining (unrewarded) bowl until the animal loses interest and the trial is ended. After correct choices, the trial ends once the animal has consumed the food reward. E. Averaged traces for ipsilateral mPFC → MD sRGECO responses to RS errors (vertical dashed line) across groups (line represents means, shaded regions represent SEM). Light gray = eYFP saline first; dark gray = eYFP psilocybin first; teal = ChR2 saline first; dark blue = ChR2 psilocybin first. F. ChR2 saline first animals have significantly higher mPFC → MD neuron activity during the post-exploration period (10-90s) after RS errors on Day 2, the same day they have a cognitive flexibility impairment. * p < 0.05, ** p < 0.01, *** p < 0.001 by repeated measures two-way ANOVA and Tukey’s post-hoc test (C, F). See also Figure S4.

### Psilocybin elicits plasticity in prefrontal subcortically-projecting neurons

Given that enhanced structural plasticity in the cortex is a well-established effect of psychedelic drugs^3,4^, we investigated whether subcortically-projecting (SC) neurons might undergo functional plasticity that could explain the protective effects of psilocybin. To measure synaptic plasticity, whole-cell recordings were performed in Layer 5 (L5) SC neurons 24 hours after the injection of 2 mg/kg psilocybin or saline (control), and MD terminals expressing ChR2 were stimulated with light (**Figure 4A**). SC neurons, known to project to MD thalamus among other subcortical regions, can be physiologically classified based on their h-current, using a previously established metric^25^: the summed amplitude of their voltage sag and rebound in response to a hyperpolarizing current pulse (**Figure S4 A-B**). Psilocybin did not alter intrinsic properties such as input resistance, resting membrane potential, amplitude of voltage sag or rebound, and time to sag peak in SC neurons (**Figure 4B-C, S5C-D**). However, we did observe a robust synaptic potentiation in cells from psilocybin-treated animals, with increased amplitude and rise slopes of optogenetic EPSPs (oEPSPs) and optogenetic EPSCs (oEPSCs) (**Figure 4D-I**). Other than a significant increase in the fast component of the decay time constant in oEPSCs recorded from psilocybin-treated mice, there were no differences in decay rate, half-width, and time to peak for oEPSCs and oEPSPs (**Figure S5E-N**). In neurons recorded from psilocybin-treated animals, we also observed increased paired-pulse depression of oEPSCs for interpulse intervals from 50-200ms (**Figure 4J-K, S5Q-R**). This suggests that increased oEPSC and oEPSP amplitude may reflect higher presynaptic release probability from a readily releasable pool that is quickly depleted.

**Figure 4.**
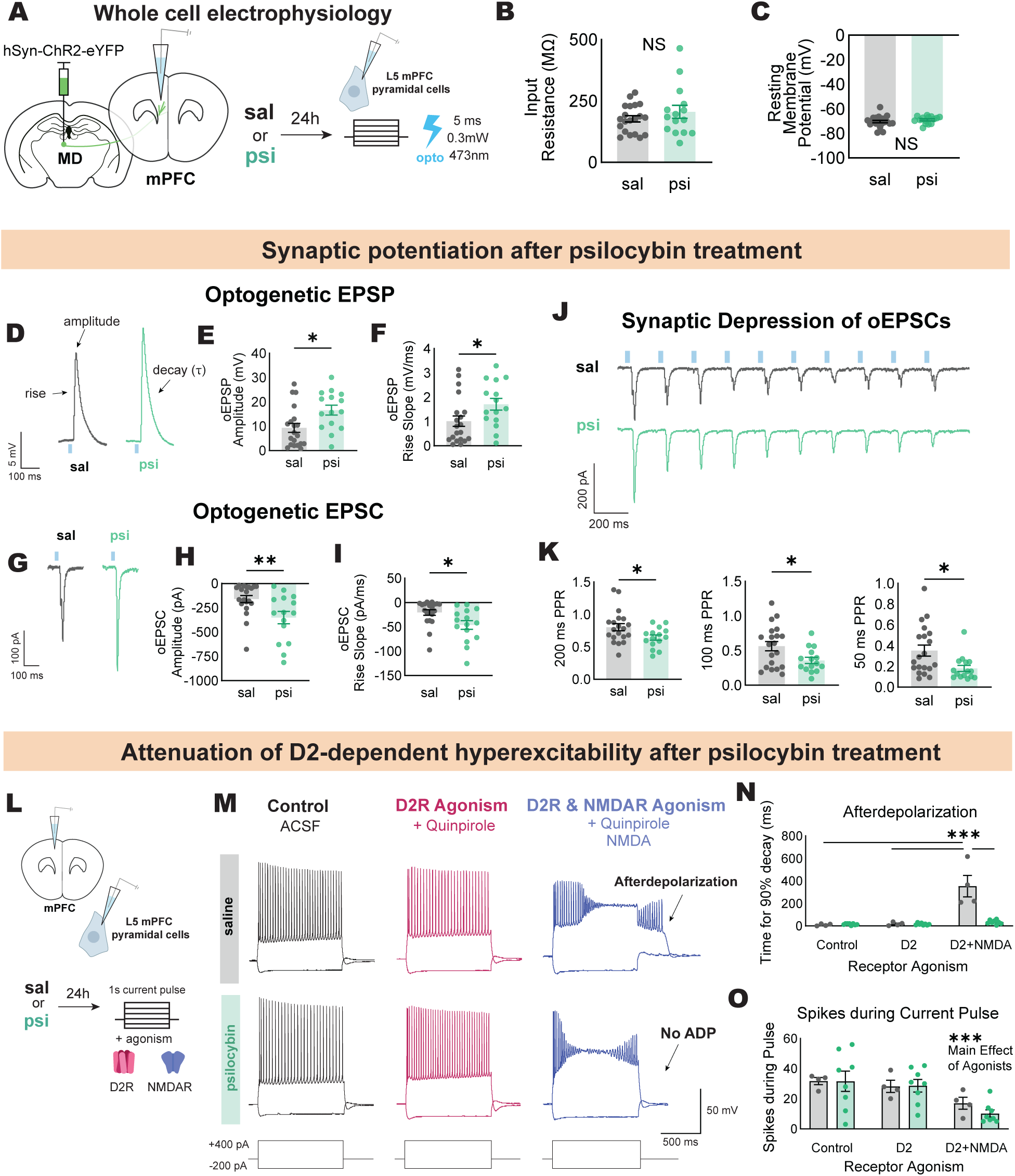
Psilocybin elicits plasticity in prefrontal subcortically-projecting neurons. A. Schematic of experimental design for whole-cell recordings of Layer 5 mPFC pyramidal cells 24 hours after psilocybin or saline injection, with optogenetic stimulation of terminals from mediodorsal thalamus (MD) inputs. B. No differences in input resistance were observed between cells from saline-treated (5 mice, 20 cells) vs. psilocybin-treated mice (4 mice, 15 cells). C. Same as B but for resting membrane potential. D. Representative sample EPSPs triggered by a 5ms pulse of 473 nm light (blue bar). E. Optogenetic EPSP (oEPSP) amplitude is significantly greater in cells from psilocybin-treated mice. F. Rise slope in the first 4ms from inflection point is significantly greater in cells from psilocybin-treated mice. G. Representative sample EPSCs triggered by a 5ms pulse of 473nm light (blue bar). H. Same as in E but for oEPSCs. I. Same as in F but for oEPSCs. J. Representative sample traces of oEPSCS recorded from a pulse train with 200 ms interpulse interval (blue bars: light pulses). K. Paired pulse ratio is significantly decreased in cells from psilocybin-treated mice whether the interpulse interval is 200 ms (left), 100 ms (center), or 50 ms (right). L. Schematic of experimental design for whole-cell recordings of Layer 5 mPFC pyramidal cells with quinpirole and NMDA to examine the effects of psilocybin or saline treatment on the quinpirole-dependent afterdepolarization. M. Representative sample traces of cellular recordings from saline-treated animals (top) or psilocybin-treated animals (bottom) during control conditions (black, left), with quinpirole only (red, center), and after wash-in of both quinpirole and NMDA (blue, right). The afterdepolarization occurs following depolarizing current pulses which evoke spiking. N. Cells recorded from saline-treated animals (2 mice, 4 cells) exhibit a significantly longer afterdepolarization compared to psilocybin (5 mice, 8 cells), quantified as the amount of time for the membrane potential to decay 90% from peak amplitude (after the end of the depolarizing current pulse). O. The number of spikes occurring during the current pulse is affected by quinpirole and NMDA based on the agonists in bath solution, but there was no significant difference between the saline and psilocybin conditions. * p < 0.05, ** p < 0.01, *** p < 0.001 by unpaired two-tailed t-test (B, C, E, F, H, I, K) or repeated measures two-way ANOVA with Tukey’s post-hoc test (N, O). Bars represent means, error bars represent SEM, and dots represent individual cells. See also Figure S5.

L5 SC neurons express dopamine D2 receptors (D2Rs) which our laboratory has previously shown to induce prolonged afterdepolarizations (ADPs) in conjunction with synaptic NMDAR activation^25,26^. These D2R-driven ADPs are well-positioned to contribute to abnormal elevations in post-error activity elicited by stimulation of TH+ VTA → mPFC projections. We therefore sought to examine the extent to which psilocybin might alter these ADPs. For this, we performed current-clamp recordings in L5 SC neurons before, during, and after bath-application of the D2R agonist quinpirole and NMDA (**Figure 4L**). Remarkably, the ability to induce an ADP was almost completely abolished in neurons from psilocybin-treated animals (**Figure 4M-N**). In the absence of NMDA, the spike rate in response to current injection was similar for neurons collected from saline- and psilocybin-treated animals (**Figure 4O**), suggesting this reflects a specific alteration of this D2R-dependent mechanism, rather than a nonspecific change in overall excitability. These experiments reveal that psilocybin can induce both synaptic and intrinsic plasticity in mPFC L5 pyramidal neurons that project subcortically, including to MD thalamus.

## DISCUSSION

Here, we demonstrate that psilocybin alters learning strategies in both healthy and impaired animals, and can restore cognitive flexibility in a manner that depends on the specific mechanism underlying the cognitive dysfunction. We further show that the optogenetic induction of cognitive inflexibility alters activity in a specific circuit known to be crucial for flexible decision-making, and that psilocybin reverses this alteration both acutely and persistently, paralleling its behavioral rescue. Finally, we identify cellular and synaptic plasticity at this locus that could explain these effects.

### Psilocybin increases flexibility in healthy and pathological states

Previous studies of psychedelic drugs in rodents have found improvement in cognitive flexibility in highly-trained, healthy animals^27–30^. First, in our naturalistic home-cage digging task, animals have minimal exposure to the cues and task mechanics before testing, requiring little to no training. Second, to our knowledge, this is the first study measuring the effects of psychedelic drugs on cognitive inflexibility induced by targeted circuit manipulations.

In this context, psilocybin increases flexibility in healthy mice, decreasing “stay” transitions and increasing “shift” transitions whether the prior outcome was a “win” or “loss”, respectively. This effect was relatively modest but confirms that psilocybin can affect flexibility even under baseline conditions and provides some information about how this might occur. In mice that receive deleterious VTA → mPFC optogenetic stimulation, psilocybin rescues cognitive inflexibility. Interestingly, inflexibility associated with VTA → mPFC optogenetic stimulation is associated with a significant increase in errors on both conflict and non-conflict trials, whereas inhibiting cPV synapses elicits minimal increase in errors on non-conflict trials. Given that psilocybin seems to enhance flexibility particularly on non-conflict trials, this may be one reason why psilocybin rescues cognitive inflexibility when it is induced by VTA → mPFC optogenetic stimulation but not when it is the result of inhibiting cPV synapses.

### PFC → MD neurons as a key locus underlying therapeutic and pathological changes in flexibility

For the first time, we identify a possible mechanism for psilocybin’s therapeutic effects on flexibility deficits. TH+ VTA → mPFC burst stimulation abnormally elevates mPFC → MD neural activity during post-error exploration. Psilocybin attenuated this elevation in mPFC → MD activity acutely and continued to do so 24 hours after administration, restoring both normal neural activity and performance in the cognitive flexibility assay. The enduring protective effects of psilocybin could be due to functional plasticity in Layer 5 (L5) pyramidal neurons that project subcortically, including to MD thalamus. Indeed, other studies have found that psilocybin induces functional plasticity in L5 pyramidal cells^4^, with long-lasting therapeutic effects involving neurons that project subcortically, including^31^ or excluding^32^ thalamus. We specifically found a potentiation of synaptic input from MD to L5 subcortically-projecting neurons. These neurons also lost their pronounced dopamine D2 receptor (D2R)-dependent afterdepolarization (ADP) after psilocybin treatment, providing a mechanism for psilocybin’s attenuation of mPFC → MD post-error activity. Through this ADP, D2Rs and synaptic input through NMDAR activation^26^ can enhance the excitability of these neurons on long timescales, even up to 10s^25^. Burst stimulation of TH+ VTA → mPFC projections, which release dopamine and glutamate^19^, may trigger this hyperexcitable state in mPFC → MD neurons, causing abnormally elevated activity during post-error exploration. By blocking this ADP, psilocybin may restore normal post-error neural signaling within the mPFC → MD population, thereby protecting against stimulation-induced behavioral deficits.

The convergence on mPFC → MD neurons is notable, because several studies from our laboratory and others have implicated this population in cognitive flexibility. We previously found that inhibiting mPFC → MD projections disrupts shifts from auditory to visual cue-based rules^18^. Another study found that disrupting mPFC → MD activity specifically during outcome periods following conflict trials is sufficient to impair learning in a similar set-shifting task, suggesting that post-error activity in this population plays a key role in learning the new rule^24^. Recent work from our laboratory has shown that callosal GABAergic synapses from parvalbumin (PV) neurons onto mPFC → MD neurons become strengthened during the learning of rule shifts^20^. We also found that gamma-frequency synchronization between prefrontal PV neurons and mPFC → MD neurons during the pre-decision period changes when mice learn the rule shift, and distinguishes between conflict and non-conflict trials^21^. Other studies have shown that prefrontal-MD interactions are important for using context to switch between the use of different sensory modalities for decision-making^33,34^.

### Dosage considerations

When psychedelic drugs are administered at sufficiently high doses, even just one or two treatments can induce long lasting improvements in mood and cognitive flexibility^9,10,35^. Therapeutic outcomes correlate with the intensity of the subjective experience, which scale with dose and occupancy of the 5HT2A serotonin receptor^36^. In humans, doses ranging from 20-30 mg/70kg have similarly strong subjective effects^37^, which correspond to 60-66% 5HT2A receptor occupancy^36^. A study in mice demonstrates that subcutaneously administered 1 mg/kg psilocybin corresponds to 44% receptor occupancy, and 3.2 mg/kg corresponds to 62% occupancy. The subjective effects of hallucinogenic drugs are inferred in rodents through the head-twitch response, which has predictive validity for the subjective potency of hallucinogens in humans^38^. Both the 1 mg/kg and 3.2 mg/kg dose induced a robust head-twitch response, but the 3.2 mg/kg dose decreased locomotion^39^. While the % 5HT2AR occupancy for 2 mg/kg dose of psilocybin used here is not directly known, this dose when administered intraperitoneally (IP) induces head-twitches without locomotor deficits^40^. The 2 mg/kg IP dose also induced acute anxiety-like behaviors in mice^40^, which may reflect the challenging subjective experience that high doses of psilocybin are known to induce^41^. Thus, a 2 mg/kg dose in mice is well-suited to mimic the high doses that cause high 5HT2AR occupancy in humans without impairing the ability to perform the behavioral tasks required to measure cognitive flexibility.

### Future directions

Direct pharmacological blockade of the D2R-dependent ADP during stimulation could determine whether this is a therapeutic mechanism that can be targeted, distinct from other actions of psychedelic treatment. A major open question is how psilocybin abolishes the ADP in L5 mPFC → MD cells, which should be examined in future work. Our results provide a cellular, circuit, and behavioral mechanism for psilocybin’s therapeutic effects that involves the midbrain, cortex, and thalamus, brain regions implicated in many psychiatric disorders. Thus, examining how psilocybin modulates functional interactions between these regions, e.g., by measuring oscillatory synchronization, could further validate the ability of psilocybin to modulate these interactions *in vivo* during specific behavioral states.

## Conclusion

The effects of psychedelic drugs are profound and only beginning to be understood. Cognitive flexibility impairments are observed across many psychiatric disorders, such as depression, schizophrenia, autism, addiction, and PTSD^13^. Deficits across these disorders may be caused by overlapping or distinct neurobiological mechanisms. There are likely many circuit mechanisms that cause cognitive dysfunction and are responsive to treatment with psychedelic drugs, while others might be unaffected or even become worse after psychedelic treatment. Importantly, our study shows that psilocybin can rescue behavioral deficits that are not simply the direct result of altered serotonin signaling. Given the immense potential for the use of psychedelic drugs in psychiatric treatment, our results provide a timely framework for a precision medicine approach to optimize therapeutic outcomes.

## RESOURCE AVAILABILITY

### Lead Contact

Further information and requests for resources and reagents should be directed to and will be fulfilled by the lead contact, Vikaas S. Sohal.

### Materials Availability

This study did not generate any new reagents.

### Data and code availability

All data and code will be shared by the lead contact upon request.

## ACKNOWLEDGEMENTS

We thank Lynnell Mak, Emmie Hou, Joseph Baran, Joey Huang, and Aspen Kim for technical assistance. This work was supported by funding from the National Institutes of Health (K12GM081266 to C.L., T34-GM145400 to I.M.L., R01MH129835 to V.S.S., and R01MH128364 to V.S.S., T. Nowakowski, and J. Rubenstein), the UC President’s Postdoctoral Fellowship Program (C.L.), the UCSF office of the Associate Vice Chancellor for Research Opportunity and Impact (to C.L.), and philanthropic funding by a donor who wishes to remain anonymous.

## AUTHOR CONTRIBUTIONS

C.L. and V.S.S. designed the experiments and analyses. C.L. performed the analyses. C.L. and E.M.H. performed behavioral experiments. A.S.E. and U.B.B. performed ex vivo electrophysiology experiments. C.L. and I.M.L. performed stereotactic surgery. C.L., E.M.H., I.M.L., and A.R.K. performed histology. E.M.H. and A.R.K. assisted in analyses. C.L. and V.S.S. drafted the manuscript. All authors reviewed the manuscript before submission.

## METHODS

### Experimental model and subject details

#### Mice

All animal care procedures and experiments were conducted in accordance with the National Institutes of Health guidelines and approved by the Administrative Panels on Laboratory Animal Care at the University of California, San Francisco. Mice were housed in a temperature-controlled environment (22–24°C) with ad libitum access to food and water until the onset of rule-shift experiments. Mice were reared in normal lighting conditions (12-h light/dark cycle) and were moved to a reverse light/dark schedule and singly housed for the duration of behavior experiments. During this time, mice were food restricted over the course of ∼1 week so that they were 80-85% of baseline weight at the start of each task session (prior to consuming food rewards) and >85% of their baseline weight after performing the task (and consuming food rewards) each day. Male and female mice (4–9 weeks old at the time of initial surgeries) from the following lines were used: C57BL/6 (JAX: 000664), PV-Cre:Ai14 (generated in-house from JAX: 008069 and 007914), TH-Cre (line FI12, GENSAT). All experiments were performed with age-matched littermates of both sexes when possible.

#### Drugs

Psilocybin (received from the NIDA Investigational Drug and Material Supply) was prepared as described previously^40^. Briefly, psilocybin was dissolved in 0.9% sterile injectable saline at a concentration of 0.2 mg/mL and delivered at a volume of 10 μL/g intraperitoneally (e.g. 0.25 mL for a 25 g mouse) for a dose of 2 mg/kg 5 minutes before behavior experiments.

#### Stereotactic Surgery

Mice were anesthetized with isoflurane and positioned in a stereotaxic frame (David Kopf Instruments), with body temperature maintained using a heating pad. After exposing the skull, stereotaxic alignment was confirmed using bregma and lambda. The bregma-lambda distance (BL) was used to normalize the anterior-posterior coordinate for VTA injections and implants. Viral injections were delivered at a rate of 100-150 nL/min using a 10 μL Nanofil syringe (World Precision Instruments) fitted with a 35-gauge beveled needle and controlled by a syringe pump (UMP3 UltraMicroPump; World Precision Instruments). Unilateral injection sides were counterbalanced across animals for behavioral experiments. The injection needle was withdrawn 5 min after the end of the infusion. Fiber-optic implants were affixed to the skull with adhesive cement (C&B Metabond; Parkell) followed by cranioplastic cement (Ortho-jet; Lang Dental). The incision was closed with tissue adhesive (Vetbond; 3M). Following surgery, mice recovered on a heated pad until ambulatory and were then returned to a clean home cage with their litter mates and provided with moistened food. Post-operative anesthesia and health checks were provided for at least 2 days after surgery. Behavioral experiments were performed after at least 6 weeks of virus expression and ex vivo electrophysiology experiments (**Figure 4 and S5**) were performed 4 weeks after virus injection. Injection sites and implant placements were confirmed in all animals by preparing coronal sections.

For experiments involving cell-body stimulation of TH+ VTA → PFC projections (**Figures 2, S2, 3, S4**), wildtype C57BL/6 mice were injected unilaterally with 150-200nL (per depth) AAVrg-rTH-Cre in the PFC (+1.7mm anterior-posterior (AP), ±0.3mm mediolateral (ML), three depths: −2.25, −2.0, −1.75mm dorsoventral (DV) and 600nL of AAV5-EF1α-DIO-hChR2(H134R)-eYFP (UNC Vector Core) or AAV5-EF1α-DIO-eYFP (UNC Vector Core) in the ipsilateral VTA (−3.4 AP, ±0.3 ML, −4.2 DV). A mono fiber-optic implant (200 μm, 0.22 NA, Doric Lenses: MFC_200/240-0.22_5.5mm_ZF1.25(G)_FLT) was inserted at a 15° angle (−3.4 AP, ±1.5 ML, −3.9 DV) to target the ipsilateral VTA. For experiments involving terminal stimulation of TH+ VTA → PFC projections (**Figure S2**), wildtype C57BL/6 or TH-Cre +/− littermates were injected with ChR2 or eYFP control at the same coordinates above, but a mono fiber-optic implant (200 μm, 0.22 NA, Doric Lenses: MFC_200/240-0.22_2.0mm_FLT) was targeted to the ipsilateral PFC (+1.7 AP, ±0.3 ML, 1-8 DV). For experiments involving cross-hemispheric inhibition of callosal parvalbumin (cPV) projections (**Figure S3**), PV-Cre:Ai14 mice were injected bilaterally in the PFC (same coordinates as above; three depths) with AAV1-CAG-DIO-Ace2N-4AA-mNeon (Virovek) and unilaterally in the PFC (+1.7 AP, ±0.3 ML, −2.5 DV) with either 1 μl of AAV2-EF1α-DIO-eNpHR3.0-BFP (Virovek) or AAV2-hSyn-BFP (Virovek). Bilateral mono fiber-optic implants (400 μm, 0.48 NA, Doric Lenses: MFC_400/430-0.48_2.8mm_ZF1.25_FLT) were positioned above the PFC (+1.7 AP, ± 0.76 ML, −1.8 DV) at a 12° angle. For experiments combining TH+ VTA → PFC cell-body stimulation with PFC → MD photometry (**Figure 3 and S4**), AAVrg-rTH-Cre and either DIO-ChR2-eYFP or DIO-eYFP control was injected unilaterally in the PFC and VTA respectively as described above, along with bilateral injections of 200 nL AAVrg-EF1α-FlpO in the mediodorsal thalamus (−1.7 AP, ± 0.35 ML, −3.7 DV). Bilateral injections of AAV8-Ef1a-Coff/Fon-sRGECO of 150-200nL per depth (three depths; same coordinates described above) and mono fiber-optic implants (400 μm, 0.48 NA, Doric Lenses: MFC_400/430-0.48_2.8mm_ZF1.25_FLT) were positioned above the PFC at a 12° angle (same coordinates described above). For ex vivo electrophysiology experiments (**Figure 4**), 4 week old mice were injected with 350 nL AAV5-EF1α-DIO-ChR2-eYFP (UNC Vector Core) in the MD (−1.7 AP, +0.3 ML, and −3.43 DV).

#### Rule-Shift Behavior

See **Figures 1A, 1C, and 3D**. Methods are adapted from the lab’s prior work^15–17^, with the modification of a barrier to separate the task and holding area within the same home-cage. Mice are first acclimated to a reverse light/dark cycle and introduced to the digging media (Sand: Mosser Lee White Sand Soil Cover, Litter: Natural Integrity Clumping Clay Cat Litter or Dr. Elsey’s Ultra Unscented Cat Litter), odor cues (Garlic or Coriander powder; McCormick), digging bowls (Nibble Bowl Small; All Living Things), and food rewards (Bio-Serv 20 mg balanced-diet pellets or Reese’s Peanut Butter Chips) until they reach 85% of their baseline weight. Mice then undergo habituation in their home cage where they are trained to dig in one of two bowls containing a compound cue consisting of an odor (Garlic or Coriander) and texture (Sand or Litter). The compound cues are organized into two presentations such that all 4 cues are present individually across the 2 compound cues (Pair 1: Coriander-scented Litter in one bowl and Garlic-scented Sand in the other; Pair 2: Garlic-scented Litter in one bowl and Coriander-scented Sand in the other). During 8-12 habituation trials, mice are allowed to dig in either bowl until they discover and consume the reward. Mice are returned to a holding area, either a separate, clean cage (experiments in **Figure 1, S1**) or a region within the home cage that is separated from the area where the next trial is performed by a barrier that is manually removed and replaced by the experimenter between trials (**Figures 2, S2, S3, 3, and S4**). During habituation, cues are evenly rewarded to ensure that mice will discover reward in each compound cue presentation and no cue-reward association is formed. Habituation typically takes 8 trials for mice to learn to dig in bowls to search for food reward, and switch to the other bowl when reward is not found. On the next day, mice perform the task where the bowl containing the food reward is determined by a specific rule (i.e. if the rule is ‘Sand’, the mouse must dig in Garlic-scented Sand when Pair 1 is presented, or in Coriander-scented Sand if Pair 2 is presented). If mice dig in the correct bowl and discover the reward, the barrier is lowered after they finish consuming the reward (keeping them in a holding area between trials) and the next trial begins in ∼30s. If the mouse chooses to dig in the incorrect bowl, the other bowl containing the reward is removed from the cage by the experimenter and the mouse is allowed to continue digging in the incorrect bowl and investigate the cage until they lose interest in searching for the reward. This self-directed post-error exploration lasts ∼150s on average. The experimenter then lowers the barrier and the next trial starts after a minimum of 90s as an error-induced time-out. After the mouse successfully learns this Initial Association (IA) by reaching 8/10 correct trials, 3 additional IA trials are performed and then the rule is shifted to the other modality (i.e. if Sand was the IA rule, the rule shift (RS) rule must be an odor cue, not a texture cue). When applicable, optogenetic manipulation begins immediately upon reaching the 8/10 IA criterion, so the 3 additional IA trials after reaching the criterion performed while the mouse is subject to the optogenetic manipulation. In experiments involving VTA → mPFC stimulation, a single unilateral burst is delivered when mice make an error during the RS stage of the task, but also if they make an error in the 3 additional trials after IA criterion is reached. For cPV inhibition experiments (**Figure S3**), the 638 nm laser is turned on after IA criterion is reached and remains on for the 3 additional IA trials and the entire duration of the RS stage. By design, the first RS trial is a Conflict Trial (see **Figure 1C**) due to the conflicting strategies to make a correct choice; following the IA rule will result in a Conflict Error. The other bowl pair presentation is considered a Non-Conflict Trial, because the correct choice during RS is the same as the correct choice during IA. When mice make 8/10 correct choices based on the RS rule, the animal reaches RS criterion and the task is concluded for that day. If mice fail to reach RS criterion by trial 40, the task is concluded and the ceiling of 40 is logged as the value for their performance that day. On all task days, animals received intraperitoneal injections of saline or 2 mg/kg psilocybin dissolved in saline 5 minutes before IA, at a volume of 10μL/g. Experiments were always performed with age-matched littermate controls and experimenters were blinded to virus and drug conditions to the best of our ability.

#### *In vivo* Optogenetic Manipulation

For behavioral experiments using optogenetic ChR2 stimulation (**Figures 2, S2, 3, S4**) A 473 nm blue laser (OEM Laser Systems) was coupled to the mono fiber-optic cannula (Doric Lenses) with a zirconia sleeve (Doric Lenses) through a 200 μm diameter mono fiber-optic patch cord (Doric Lenses) and adjusted such that the final light power was ∼5 mW. A custom Arduino-based stimulator delivered a 50 Hz pulse train of 25 4ms pulses (total duration 500ms) triggered by the experimenter during RS errors as the experimenter removed the bowl containing the reward. For optogenetic inhibition of cPV terminals (**Figure S3**), a 638nm laser (Doric Lenses) was adjusted to deliver ∼2.5mW continuously after animals reached IA criterion.

### TEMPO Recordings of Voltage Indicators

#### Optical apparatus

Methods for TEMPO recording of cross-hemispheric synchrony are described in detail in prior work^16,17,21^. Mono fiber-optic implants (400 μm core, NA = 0.48, low-autofluorescence fiber; Doric Lenses, MFC_400/430-0.48_2.8mm_ZF1.25_FLT) were stereotaxically implanted bilaterally in PFC. A matching fiber-optic patch cord (Doric Lenses, MFP_400/430/1100-0.48_2m_FC-ZF1.25) connected the implant to an optical ‘mini-cube’ (Doric Lenses, FMC5_E1(460-490)_F1(500-540)_E2(555-570)_F2(580-600)_S). The fiber on the ipsilateral side of the viral injection of either 1 μl of AAV2-EF1α-DIO-eNpHR3.0-BFP (Virovek) or 1 μl of AAV2-hSyn-BFP (Virovek) was used to both deliver excitation light to and collect emitted fluorescence from that recording site. The fiber contralateral to the viral injection site was connected to a separate mini-cube (Doric Lenses, FMC6_E1(460-490)_F1(500-540)_E2(555-570)_F2(580-600)_O(628-642)_S) that was attached to a 638 nm laser (Doric Lenses) and was used to deliver excitation light and optogenetic inhibition to and collect emitted fluorescence from that recording site. Laser power was set to 2.5mW and measured using a ThorLabs light meter (PM100D) at the fiber tip. Each 1.25 mm diameter zirconia optical implant ferrule was cleaned with 70% ethanol before each recording, then securely attached to the patch cord via a zirconia sleeve.

Excitation light for the two color channels were provided by fiber-coupled LEDs (Thorlabs M490F3 and M565F3) connected to the mini-cube by a patch cord (200 μm, NA = 0.39; Thorlabs M75L01). LEDs were controlled by a 4-channel, 10-kHz-bandwidth current source (Thorlabs DC4104). LED current was adjusted to give a final light power at the animal (averaged during modulation, see below) of approximately 200 μW for the mNeon channel (460–490 nm excitation), and 100 μW for the Red channel (555–570 nm excitation). Each of the two emission ports on the mini-cube was connected to an adjustable-gain photoreceiver (Femto, OE-200-Si-FC; Bandwidth set to 7 kHz, AC-coupled, ‘low’ gain of ∼5 × 107 V W−1) using a large-core high-NA fiber to maximize throughput (600 μm core, NA = 0.48 (Doric lenses, MFP_600/630/LWMJ-0.48_0.5m_FC-FC). All four signals (green and red, bilaterally) are then demodulated and then digitized by a multichannel real-time signal processor (Tucker-Davis Technologies, RX-8). The commercial software Synapse (Tucker-Davis Technologies) was used for data acquisition.

#### Analysis of fiberoptic TEMPO data

Interhemispheric synchrony of oscillations in the gamma-band frequency (30-50 Hz) were analyzed as described in prior work^21^. Briefly, signals acquired from the Synapse software were band-pass filtered to isolate signals in the 30-50 Hz gamma frequency band. Signal cleaning was performed via linear regression between the green Ace2N-mNeon voltage sensor channel and the red tdTomato control signal and subtraction of shared noise from the Ace2N-mNeon signal to obtain a voltage signal corrected for shared non-voltage-related changes in fluorescence. The peak phase-lag was quantified for each 250 ms bin for the full recording by calculating Pearson correlation coefficients between signals collected from the left and right hemispheres, shifting one signal relative to the other in 22.5° increments. Peak-zero-phase-lag gamma synchrony occurred when the correlation coefficient was highest for that 250 ms bin when the signals were not phase-shifted. Post-error synchrony (**Figure S2D**) was calculated by averaging the correlation coefficients of 250 ms bins with peak-zero-phase-lag synchrony from 0-30s post-error, beginning when the correct bowl was removed by the experimenter after mice dug in the incorrect bowl.

### Photometry Recordings of Calcium Indicators

#### Optical apparatus

Bulk photometry recordings of a calcium sensor were performed as described previously^16^, with some modifications to allow dual-site recordings of a red calcium sensor. Briefly, mono fiber-optic implants (MFC_400/430-0.48_2.8mm_ZF1.25_FLT) were implanted bilaterally above the PFC and a matching patch cord (MFP_400/430/1100–0.48_2m_FC-ZF1.25) provided a light path between the animal and an optical “mini-cube” (FMC6_IE(400-410)_E1(460-490)_F1(500-540)_E2(555-570)_F2(580-680)_S). Two excitation LEDs (M405FP1 and M565F3; ThorLabs) were connected to the mini-cubes by patch cords (200 μm core, NA = 0.39, Doric Lenses) and controlled by an LED driver (ThorLabs DC4104) and connected to an RX-8 real-time processor (Tucker-Davis Technologies). Excitation light is delivered at 565 nm to stimulate sRGECO fluorescence in a Ca2+-dependent manner and at 405 nm for excitation of sRGECO at its isosbestic wavelength. Excitation wavelengths were sinusoidally-modulated at non-resonant frequencies (Left 565: 210 Hz, Left 405: 330 Hz, Right 565: 280 Hz, Right 405: 305 Hz). Emission signals were collected from the mini-cubes through patch cords (Doric Lenses, MFP_600/630/LWMJ-0.48_0.5m_FC-FC) and focused onto photoreceivers (Right: DFD_FOA_FC, Left: NPM_2151_FOA_FC; Doric Lenses). Modulated signals generated by the LEDs were recovered and demodulated with the RX-8 real-time processor and Synapse software. Before each recording, the 1.25 mm diameter zirconia optical implant ferrules were cleaned with ethanol and securely attached to the patch cord via a zirconia sleeve.

#### Analysis

Demodulated signals obtained from the Tucker-Davis Technologies software for 4 channels (Left 565, Left 405, Right 565, Right 405) were first truncated to omit the initial 180s (to account for large signal fluctuations from turning LEDs on and initial bleaching). A low-pass Butterworth filter at 10 Hz was applied before normalizing the 405 nm signal to the 565 nm signal with a least-squares first order polynomial fit (Python numpy polyfit function) and scaled with the slope and intercept. The dF/F for the entire recording was calculated by subtracting the scaled isosbestic signal from the signal channel before dividing by the scaled isosbestic signal (dF/F = (565nm - fitted 405nm) / fitted 405 nm). Post-outcome mean z values were calculated by averaging the Zscore of the dF/F relative to a baseline duration. For 0-5s immediate post-outcome responses, the baseline was −3 to 0s before the outcome. For +10-90s post-error exploration responses, the baseline was −3 to +10s from the outcome, to include the immediate error response in the baseline for normalizing the extended post-error response.

### Slice Electrophysiology

#### Slice preparation

Slice preparation was performed as described previously^20^. Mice were injected with 2 mg/kg psilocybin (dissolved in 0.9% sterile saline) or saline control at a volume of 10 μL/g 24 hours before slice preparation. In slice experiments with optogenetic stimulation (**Figures 4A-G and S5A-R**), experiments were performed 4 weeks after virus injection. Cutting solution was chilled and contained (in mM): 210 Sucrose, 1.25 NaH2PO4, 25 NaHCO3, 0.5 CaCl2, 7 MgCl2, 7 Dextrose, 1.3 Ascorbic Acid, 3 Na Pyruvate, 2.5 KCl. The external ACSF used for holding slices contained (in mM): 125 NaCl, 2.5 KCl, 1.25 NaH2PO4, 25 NaHCO3, 2 CaCl2, 2 MgCl2, 10 Dextrose, 1.3 Ascorbic Acid, 3 Na Pyruvate. Slices were incubated in the holding solution at 30-33° for 30 minutes and at room temperature for at least 15 minutes before recording. The recording ACSF was warmed at 32.5° and contained (in mM): 125 NaCl, 3 KCl, 1.25 NaH2PO4, 25 NaHCO3, 2 CaCL2, 1 MgCl2, 10 Dextrose.

#### Intracellular recordings

Whole-cell recordings were performed from visually-identified pyramidal cells in layer V of infralimbic or prelimbic cortex using an upright microscope (BX51WI; Olympus). Recordings were made using a Multiclamp 700A (Molecular Devices). Patch electrodes (tip resistance = 2–6 MOhms) were filled with the following (in mM): 130 potassium gluconate, 10 KCl, 10 HEPES, 10 EGTA, 2 MgCl2, 2 MgATP, and 0.3 NaGTP (pH adjusted to 7.3 with KOH). All recordings were at 32.5 ± 1 °C. Series resistance was usually 10–20 MΩ, and experiments were discontinued above 30 MΩ.

#### Optogenetically-evoked EPSP and EPSC recordings

For experiments in corticothalamic cells with optogenetic excitation (**Figure 4A-K,S5A-R**), intrinsic properties were calculated based on the current-clamp responses to a series of 600 ms current pulse injections from −250 to 230 pA (20 pA per increment). Cells were characterized as subcortically-projecting (SC) or intratelencephalically-projecting (IT) based on their voltage sag and rebound responses when hyperpolarized with a 600 ms current pulse at −230pA. Stimulation of MD → PFC terminals was performed with 5 ms flashes of light generated by a Lambda DG-4 high-speed optical switch with a 300W Xenon lamp, and an excitation filter set centered around 470 nm, delivered to the slice through a 40× objective (Olympus). Illumination was delivered across a full high-power (40×) field at 0.3 mW.

#### Bath-applied agonists

For experiments involving bath-applied agonists (**Figure 4H-K**), intrinsic properties were calculated based on current-clamp responses to a series of 1000 ms current pulse injections from −200 to +400 pA and cells were confirmed to be CT cells based on their voltage sag and rebound responses when hyperpolarized at −200 pA. ACSF containing 15 μM (−)quinpirole was bath-applied for 25 minutes to obtain current-clamp recordings of the same cell with D2R agonism. Finally, ACSF containing 15 μM (−)quinpirole and 6 μM NMDA was bath-applied for 15 minutes to obtain current-clamp recordings of the same cell during D2R and NMDAR agonism.

### Immunohistochemistry and microscopy

Histological verification of implant location and viral spread was performed as described previously^42^. All mice used for behavioral experiments were anaesthetized with an intraperitoneal injection of Euthasol and transcardially perfused with 4% paraformaldahyde in PBS. Brains were extracted and stored in 4% paraformaldahyde for 24 h at 4℃ before being stored in PBS. 50µm thick slices were obtained on a Leica VT 1000S or Leica VT1200S.

For experiments targeting TH+ VTA → PFC projections, VTA slices were incubated with primary antibody solution (rabbit anti-TH 1:1000, chicken anti-GFP 1:500) overnight followed by 2 hour incubation of secondary antibody (anti-rabbit 555 1:750, anti-chicken 488 1:400) to validate expression of virus in TH+ cell bodies. Terminal expression of virus in PFC was validated with primary (chicken anti-GFP 1:500) and secondary (anti-chicken 488 1:400) antibody solutions to amplify eYFP. For experiments targeting xPV terminals, PFC slices were incubated in primary (rabbit anti-GFP 1:500) and secondary (goat anti-rabbit 647 1:400) to amplify BFP. Slices were mounted to slides with a mounting medium containing DAPI (Vectashield or Southern Biotech). Imaging was performed using a Keyence BZ-X All-in-One Fluorescence Microscope.

### Data Analyses and Statistics

Statistical analyses were performed using GraphPad Prism 11 and detailed in the corresponding figure legends. Quantitative data are expressed as the mean and error bars or shaded areas represent the standard error of the mean (SEM). Comparison of means were performed using unpaired two-tailed Student’s t-test, two-way ANOVA (with or without repeated measures) with Tukey’s post-hoc test. Sample sizes and statistical tests are detailed in figure legends. Data are represented as mean ± SEM unless otherwise stated.

---

**Figure S1. Related to Figure 1**

A. Raw number of total RS Errors per day and drug treatment for mice receiving saline (n = 7) or psilocybin (n = 7).

B. Raw number of non-conflict RS Errors per day and drug treatment.

C. % of total RS errors that were of the non-conflict type across days and drug treatments.

D. Same as B, but for conflict Errors.

E. Same as C, but for conflict Errors.

F. Trials to reach 8/10 criterion for IA and RS across sexes, days, and drug treatments (saline female n = 3, saline male n = 4, psilocybin female n = 3, psilocybin male n = 4).

G. Change in number of non-conflict errors split by sex and drug treatment.

H. Same as in G but for conflict errors

I. Change in % non-conflict Win-Stay transitions split by sex and drug treatment.

J. Same as in I but for non-conflict Win-Shift transitions.

K. Same as in I but for non-conflict Lose-Stay transitions.

L. Same as in I but for non-conflict Lose-Shift transitions.

M. Same as in I but for conflict Win-Stay transitions.

N. Same as in I but for conflict Win-Shift transitions.

O. Same as in I but for conflict Lose-Stay transitions.

P. Same as in I but for conflict Lose-Shift transitions.

*p < 0.05, by two-way repeated measures ANOVA (A-P). Bars represent means and lines represent individual animals (A-F). Bars represent means, error bars represent SEM, and dots represent individual animals (G-P).

**Fig S2. Related to Figure 2**.

A. Schematic for stimulation of TH+ VTA → mPFC terminals in mPFC using TH-Cre+/−mice (left). AAV5-DIO-ChR2-eYFP (n = 3 mice) or AAV5-DIO-eYFP control (n = 1 mouse) was injected in the VTA and a fiber implant positioned unilaterally above the mPFC. Right: representative histological images. Scale bar = 100 microns.

B. Schematic for stimulation of TH+ VTA → mPFC terminals in mPFC using wildtype (WT) mice (left). AAVrg-rTH-Cre was injected unilaterally in the mPFC for retrograde expression of Cre in TH+ cells. AAV5-ChR2-eYFP (n = 4 mice) or AAV5-DIO-eYFP control (n = 2 mice) was injected ipsilaterally in the VTA and an optical fiber implant positioned above the mPFC. Right: representative histological images.

C. Experimental timeline: Day 1: Baseline. Day 2: a single burst of unilateral optogenetic stimulation was delivered on RS error trials.

D. Overlaid schematics for histology from individual animal showing implant locations and viral spread.

E. No difference was observed between eYFP and ChR2 mice on their performance in IA on Day 2 relative to Day 1 (empty circles: TH-Cre +/− genotype, black circles: WT littermates with AAVrg-rTH-Cre).

F. ChR2 mice required significantly more trials to reach RS criterion on the stimulation day relative to baseline, compared to eYFP controls.

G. ChR2 mice made significantly more conflict errors on the stimulation day relative to baseline, compared to eYFP controls.

H. Same as in G but for non-conflict errors.

I. Overlaid schematics for histology from individual animals showing implant locations and viral spread for eYFP mice (left) and ChR2 mice (right) used for experiments in Figure 2 (eYFP n = 21, ChR2 n = 28).

* p < 0.05, ** p < 0.01 by unpaired two-tailed t-test (C-F). Bars represent means, error bars represent SEM, and dots represent individual animals.

**Fig S3. Psilocybin fails to rescue cognitive flexibility impaired by inhibition of callosal parvalbumin projections in mPFC**

A. Schematic for optogenetic inhibition of callosal parvalbumin projections (cPV) in mPFC, combined with bilateral photometry recordings from the genetically-encoded voltage indicator Ace2N-mNeon (left). PV-Cre; Ai14 animals were injected bilaterally in the mPFC with AAV-DIO-Ace2N-mNeon and unilaterally with AAV-DIO-eNpHR-BFP or AAV-hSyn-BFP control. Bilateral implants were positioned above mPFC, and 638 nm 2.5mW continuous light was delivered to the hemisphere contralateral from the eNpHR/BFP injection to inhibit PV+ terminals on Day 2 of the experimental timeline. Right: representative histological images.

B. Overlaid schematics for individual animals showing implant locations and viral spread.

C. Experimental timeline: Day 1: Baseline (no optogenetic or drug manipulations). Optogenetic inhibition is delivered on Day 2. Drug treatment occurs on Day 4.

D. Interhemispheric mPFC synchrony in the gamma frequency (30-50 Hz) band was decreased in eNpHR mice (n = 17) compared to BFP controls (n = 12) on the day of cPV optogenetic inhibition. Dots represent individual animals, horizontal lines represent means, and error bars represent SEM.

E. Left: No differences were observed in trials required to reach IA criterion across days for BFP (control) mice that received saline (n = 6), BFP mice that received psilocybin (n = 6), eNpHR mice that received saline (n = 8), or eNpHR mice that received psilocybin (n = 9). Right: A main effect of Group was observed, with eNpHR mice requiring more trials to reach RS criterion relative to baseline, compared to BFP controls.

F. A main effect of Group was observed, with eNpHR mice making more conflict errors relative to baseline compared to BFP controls.

G. Under the acute effects of psilocybin, eNpHR mice made significantly more non-conflict errors relative to baseline compared to other days and groups.

H. No significant differences were observed across days or groups for change in % non-conflict Win-Stay transitions relative to baseline.

I. The change in % non-conflict Win-Shift transitions (relative to baseline) was higher in eNpHR mice during acute psilocybin administration compared to other task days in this group.

J. Same as in H but for non-conflict Lose-Stay transitions.

K. Same as in H but for non-conflict Lose-Shift transitions.

L. Same as in H but for conflict Win-Stay transitions.

M. Same as in H but for conflict Win-Shift transitions.

N. A main effect of Day was observed for the change in % conflict Lose-Stay transitions.

O. Same as in H but for conflict Lose-Shift transitions.

* p < 0.05, * p < 0.01, *** p < 0.001 by unpaired two-tailed t-test (D) or repeated measures two-way ANOVA with Tukey’s post-hoc test (E-O). Bars represent means and lines represent individual animals (E-O).

**Figure S4. Related to Figure 3**.

A. Overlaid histology schematics for individual animals showing implant location and viral spread.

B. No differences were observed for trials to reach IA criterion across days and groups. Dots represent individual animals.

C. Representative averaged peri-outcome traces from Baseline Day 1 for correct (yellow) and incorrect (orange) choices (top left). mPFC → MD activity immediately after outcomes (0-5s) is higher for errors than correct outcomes (top right). Yellow dots (correct) and orange triangles (error) represent the averaged responses per animal on Day 1. Schematic showing that the experimenter is removing the bowl during the immediate post-error timepoint whereas the animal is allowed to explore without experimenter interference after bowl removal (bottom).

D. No differences were observed across groups and days for the immediate response to error outcomes.

E. Averaged traces for ipsilateral PFC → MD sRGECO responses to IA errors (vertical dashed line) across groups (line represents means, shaded regions represent SEM). Light gray = eYFP saline first; dark gray = eYFP psilocybin first; teal = ChR2 saline first; dark blue = ChR2 psilocybin first.

F. Above: Averages of immediate (0-5s) response to IA Errors from trials across groups and days. Below: Same as above but for 10-90s after the IA Error.

G. Same as in E but for IA Correct outcomes.

H. No significant differences were observed for the response to IA correct outcomes across groups and days.

I. Same as in E but for RS Correct outcomes.

J. A main effect of Group was observed for the response to RS Correct outcomes.

* p < 0.05,** p < 0.01, *** p < 0.001 by repeated measures two-way ANOVA (B, C) or two-way ANOVA with Tukey’s post-hoc test (D, F, H, J). Dots represent means across trials, error bars represent SEM (D, F, H, J).

**Fig S5. Related to Figure 4**.

A. Representative sample traces for subcortically-projecting (SC) and intratelencephalically-projecting (IT) L5 pyramidal cells in response to hyperpolarizing and depolarizing current pulses.

B. SC (n = 35) vs. IT neurons (n = 5) are classified based on the summed amplitude of their voltage sag and rebound in response to a hyperpolarizing current pulse, which reflects the h-current.

C. Left: No significant differences were observed for this measure of h-current in SC neurons from saline-treated (5 mice, 20 cells) vs. psilocybin-treated mice (4 mice, 15 cells). Right: No significant differences were observed for the h-current in IT neurons from saline-treated (1 mouse, 1 cell) vs. psilocybin-treated mice (2 mice, 4 cells).

D. No significant difference was observed in the time to reach peak amplitude of voltage sag from the onset of the current pulse in SC neurons recorded from mice treated with saline vs. psilocybin.

E. No significant difference was observed in the slow decay time constant (tau; ô) in oEPSCs recorded from saline- vs. psilocybin-treated mice.

F. SC neurons recorded from psilocybin-treated mice had a significantly higher fast tau compared to saline controls.

G. Same as in E but for weighted tau.

H. Same as in E but for half-width.

I. Same as in E but for time to peak.

J. Same as in E but for oEPSPs.

K. Same as in E but for fast tau of oEPSPs.

L. Same as in E but for weighted tau for oEPSPs.

M. Same as in E but for oEPSP half-width.

N. Same as in E but for time to peak for oEPSPs.

O. Paired pulse ratio across interpulse intervals for cells recorded from saline- or psilocybin-treated mice. Main effects of Drug and Interpulse Interval were observed.

P. A significant interaction for oEPSC amplitude was observed between drug and pulse number in a 5 Hz pulse train.

Q. Same as in R but for a 10 Hz pulse train.

R. Same as in R but for a 20 Hz pulse train.

* p < 0.05, ** p < 0.01, by unpaired two-tailed t-test (B-N) or repeated measures two-way ANOVA with Tukey’s post-hoc test (O-R). Bars represent means, error bars represent SEM, and dots represent individual cells (B-O). Dots represent means and shaded regions represent SEM (P-R).

