## Supplementary figures and images for "Psilocybin selectively rescues cognitive flexibility impairments caused by aberrant prefrontal error signaling"

### Supplemental Figure 1

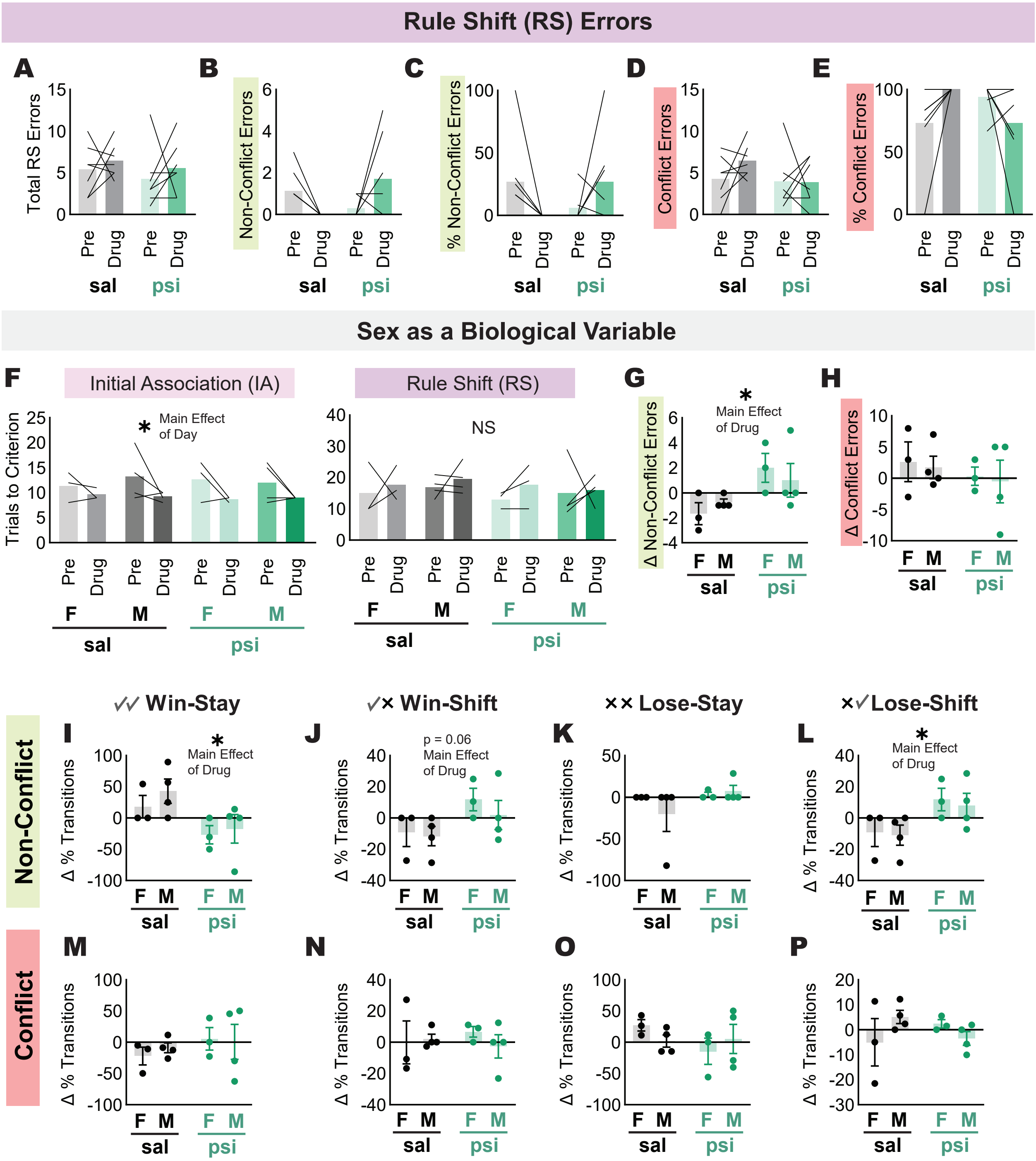

### Supplemental Figure 2

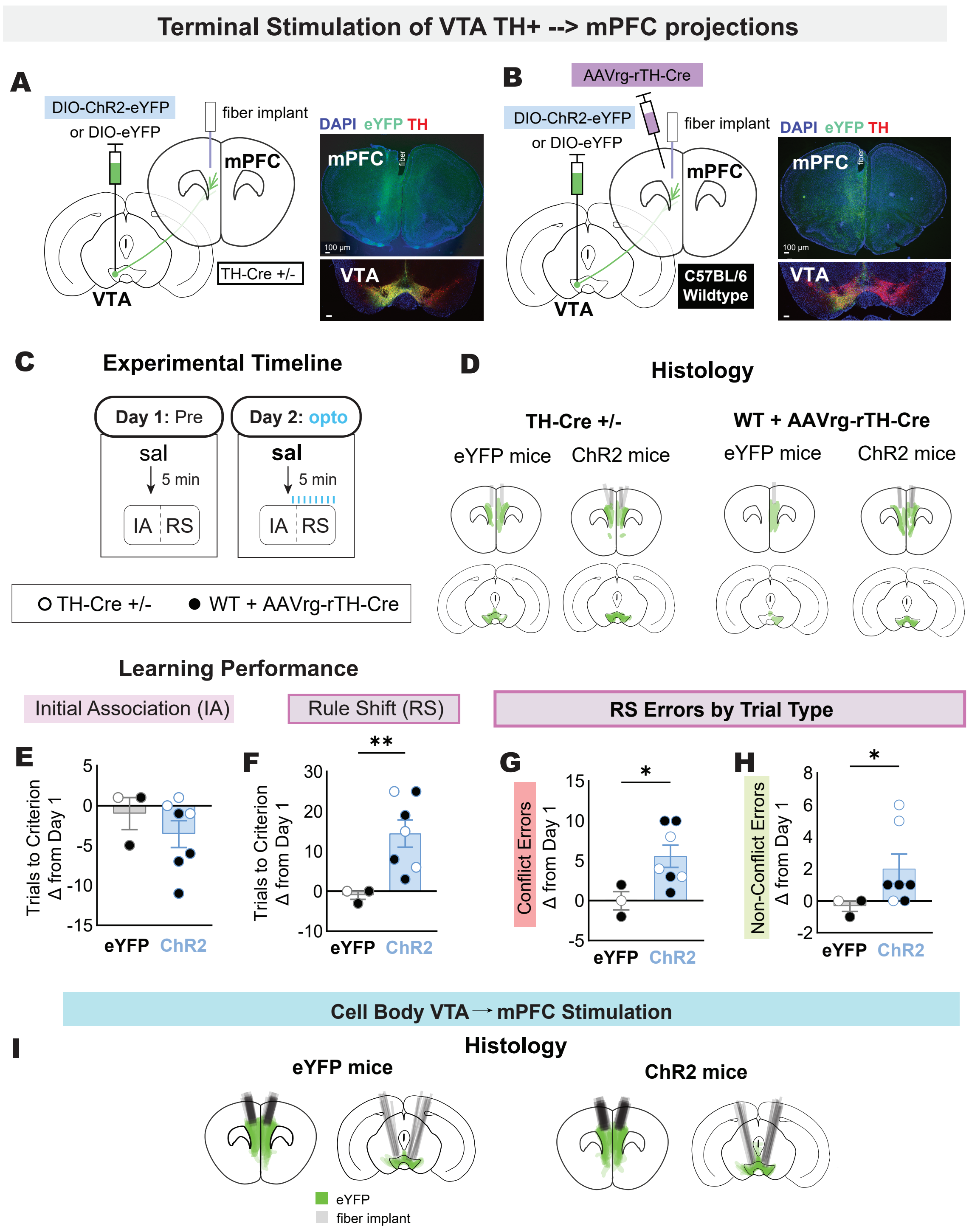

### Supplemental Figure 3

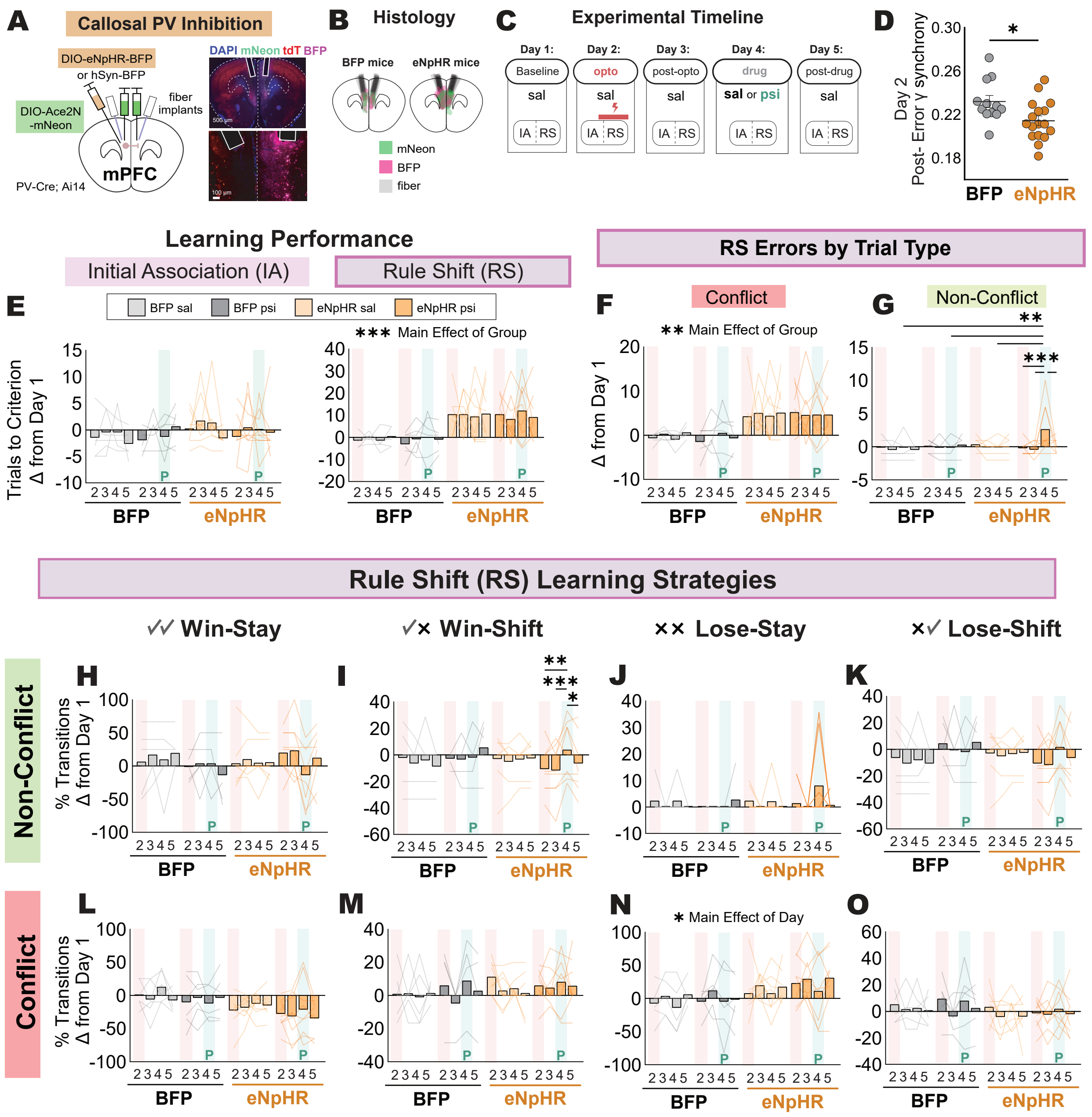

### Supplemental Figure 4

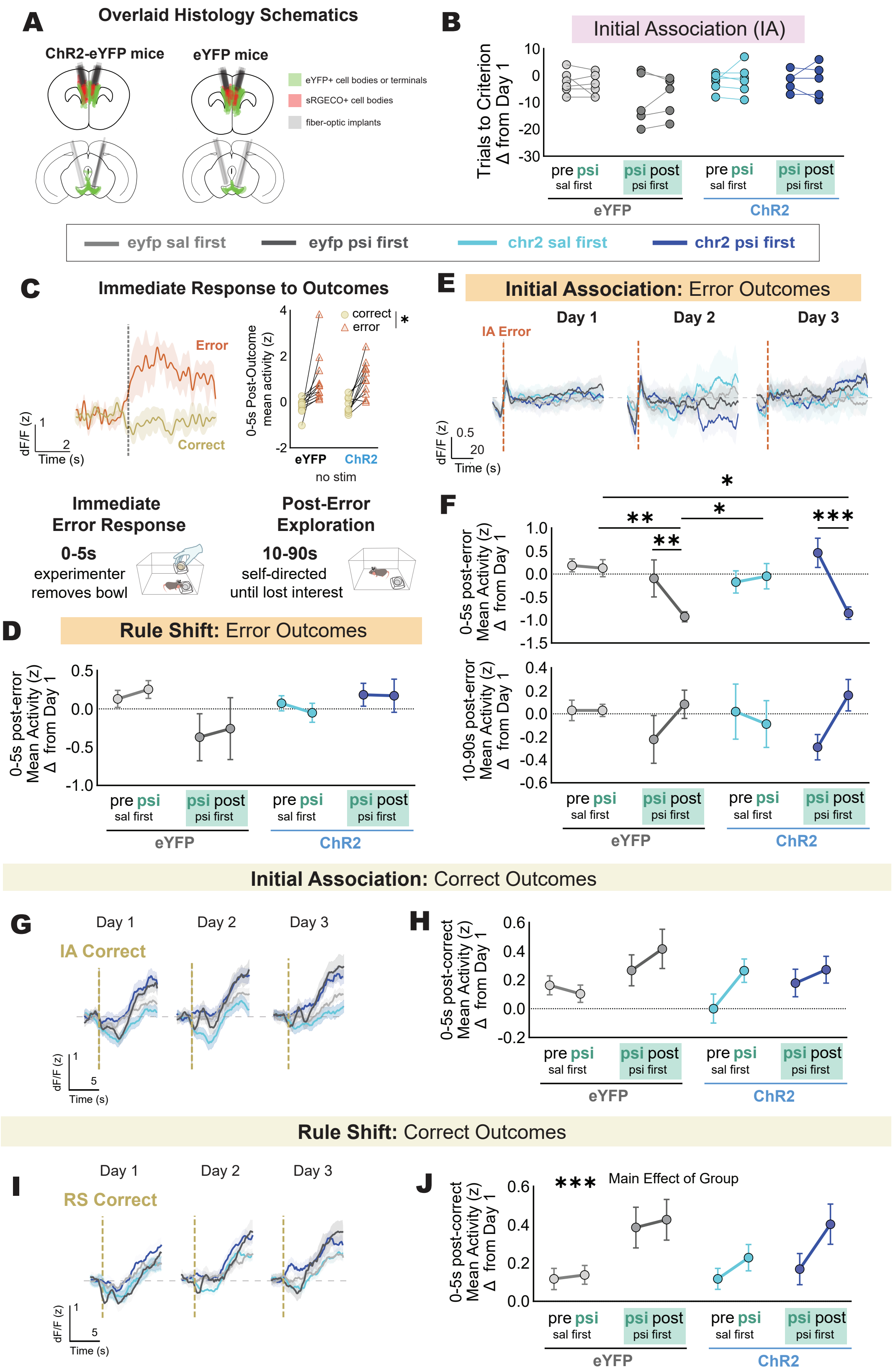

### Supplemental Figure 5

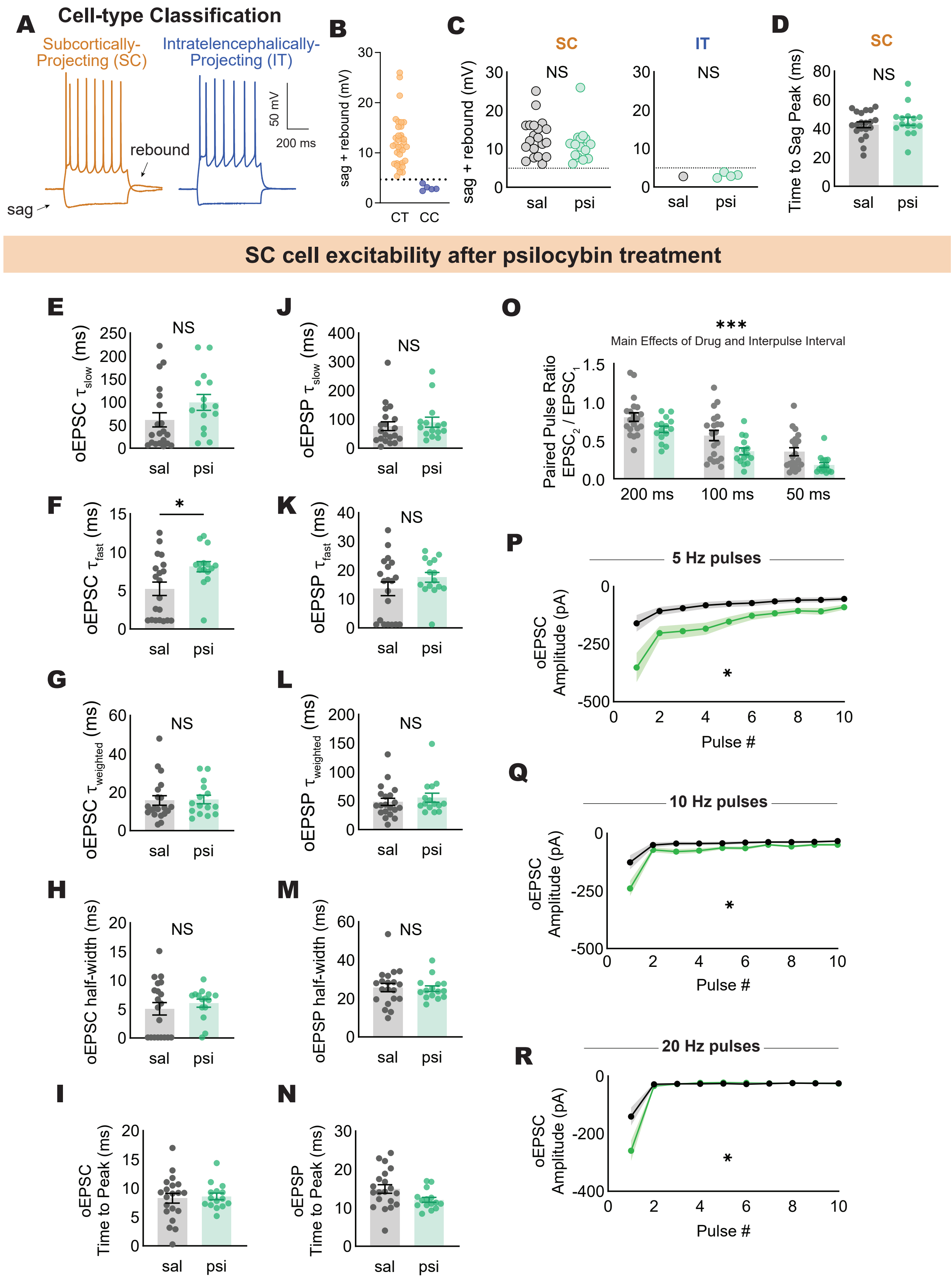
